# Basolateral amygdala CB1 receptors modulate HPA axis activation and context-cocaine memory strength during reconsolidation

**DOI:** 10.1101/2020.07.18.209932

**Authors:** Jessica A. Higginbotham, Nicole M. Jones, Rong Wang, Ryan J. McLaughlin, Rita A. Fuchs

**Author notes:** Corresponding Author: Rita A. Fuchs, PhD, Department of Integrative Physiology and Neuroscience, Washington State University College of Veterinary Medicine, PO Box 647620, Pullman, WA 99164-7620.

## Abstract

Re-exposure to a cocaine-associated context triggers craving and relapse through the retrieval of salient context-drug memories. Upon retrieval, context-drug memories become labile and temporarily sensitive to modification before they are reconsolidated into long-term memory stores. Cannabinoid type 1 receptor (**CB1R**) signaling is necessary for cocaine-memory reconsolidation and associated glutamatergic plasticity in the basolateral amygdala (**BLA**); however, it remains unclear whether CB1Rs in the BLA mediate this phenomenon. To investigate this question, we examined whether CB1R antagonist or agonist administration into the BLA immediately after cocaine-memory retrieval (i.e., during memory reconsolidation) alters cocaine-memory strength and subsequent drug context-induced cocaine-seeking behavior in an instrumental rodent model of cocaine relapse. Intra-BLA administration of the CB1R antagonist, AM251 (0.3 µg/hemisphere) – during, but not after, memory reconsolidation – increased drug context-induced cocaine-seeking behavior three days later, while the CB1R agonist, WIN55,212-2 (0.5 µg/hemisphere) failed to alter this behavior. Furthermore, AM251 administration into the posterior caudate putamen (anatomical control region) during memory reconsolidation did not alter subsequent context-induced cocaine-seeking behavior. In a follow-up experiment, cocaine-memory retrieval elicited robust hypothalamic-pituitary-adrenal axis activation, as indicated by an increase in blood serum corticosterone concentration, and this response was selectively extended by intra-BLA AM251 administration during the putative time of memory reconsolidation relative to all control conditions. Together, these findings suggest that CB1R populations in the BLA gate memory strength or interfere with memory maintenance, possibly by diminishing the impact of cue-induced arousal on the integrity of the reconsolidating memory trace or on the efficiency of the memory reconsolidation process.

## INTRODUCTION

Exposure to drug-associated environmental stimuli precipitates the retrieval of context-drug memories, thereby eliciting drug craving and relapse [1-3]. Upon retrieval from long-term memory stores, context-drug memories can become temporarily unstable and susceptible to modification. The maintenance of such labile memories requires their reconsolidation into long-term memory stores through a process that involves *de novo* protein synthesis [4] and synaptic plasticity [5]. Importantly, pathological memory reconsolidation may result in overly salient or intrusive drug memories, contributing to the etiology of substance use disorders (**SUDs**), and memory reconsolidation can be manipulated therapeutically to reduce the strength of drug memories and thus the propensity for drug relapse [6]. Therefore, elucidating the neurobiological underpinnings of cocaine-memory reconsolidation is important from a SUD treatment perspective.

Cannabinoid type 1 receptor (**CB1R**) signaling plays a critical role in cocaine-memory reconsolidation. Specifically, our laboratory has shown that systemic CB1R antagonist administration during cocaine-memory reconsolidation attenuates subsequent drug context-induced cocaine-seeking behavior [7]. Moreover, systemic CB1R antagonist administration interferes with glutamatergic transmission in the basolateral amygdala (**BLA**) [7], a site of protein synthesis-dependent memory reconsolidation [8-13]. However, questions remain about the contribution of BLA CB1R populations, because previous research indicates that BLA CB1R agonism [14] or antagonism [15] can similarly impair fear-memory reconsolidation. Furthermore, the role of BLA CB1Rs in appetitive memory reconsolidation has not been investigated.

The present study examined whether BLA CB1R signaling is necessary for cocaine-memory reconsolidation. First, we evaluated the effects of intra-BLA CB1R antagonist and agonist treatments administered immediately after cocaine-memory retrieval (i.e., during the putative time of memory reconsolidation) on memory strength, as indicated by the magnitude of subsequent context-induced cocaine-seeking behavior. Next, we examined whether the effects on memory reconsolidation were anatomically specific to CB1Rs in the BLA by manipulating CB1Rs in the adjacent posterior caudate putamen (**pCPu**). Finally, as a step toward identifying a mechanism by which BLA CB1Rs regulate cocaine-memory reconsolidation, we assessed the effects of intra-BLA CB1R antagonist treatment on blood serum corticosterone concentrations, an index of hypothalamic pituitary adrenal (**HPA**) axis activity. Our previous findings indicate that cocaine-memory reconsolidation is associated with increased HPA axis activity [16]. Furthermore, stressor-induced suppression of endocannabinoid signaling in the BLA is critical for stress-induced HPA axis activation [17]. Therefore, we predicted that intra-BLA CB1R antagonist treatment would selectively potentiate increases in corticosterone concentrations during cocaine-memory reconsolidation.

## MATERIALS AND METHODS

### Animals

Male Sprague-Dawley rats (*N*=112; 275-300 g upon arrival; Envigo Laboratories, South Kent, WA) were housed individually in a temperature- and humidity-controlled vivarium on a reversed light/dark cycle (lights on at 6:00 am). Rats received *ad libitum* access to water and 20-25 g of standard rat chow per day. Animal housing and care followed the guidelines defined in the *Guide for the Care and Use of Laboratory Animals* [18] and was approved by the Washington State University Institutional Animal Care and Use Committee.

### Food Training

To facilitate the acquisition of lever pressing for un-signaled cocaine infusions, rats were first trained to lever press for food reinforcement in standard operant conditioning chambers (Coulbourn Instruments, Holliston, MA) during a 16-h overnight session as described previously [8]. Food training was conducted in a dedicated chamber without exposure to contextual stimuli used for subsequent cocaine conditioning.

### Surgery

Twenty-four h after food training, rats were fully anesthetized using ketamine hydrochloride and xylazine (100 and 5 mg/kg, i.p., respectively; Dechara Veterinary Products, Overland Park, KS and Akorn, Lake Forest, IL). Jugular catheters were implanted into the right jugular vein. Stainless-steel guide cannulae (26-Ga, P1 Technologies, Roanoke, VA) were aimed at the BLA (−2.7 mm AP, ±5.0 mm ML, -6.6 mm DV relative to bregma) or pCPu (−2.7 mm AP, ±5.0 mm ML, -4.5 mm DV relative to bregma). Stainless-steel screws and dental acrylic anchored the cannulae to the skull. Rats received the analgesic, carprofen (5 mg/kg per day, p.o.; ClearH2O, Westbrook, ME) for 24 h before and 48 h after surgery. The catheters were maintained and periodically tested for patency, as previously described [19]. Rats received five days for post-surgical recovery.

### Cocaine Self-Administration and Extinction Training

Rats were randomly assigned to one of two distinctly different environmental contexts (**Table S1**) for cocaine self-administration training [8]. Training in the designated context was conducted for two h each day, during the rats’ dark cycle. During the sessions, active-lever responses were reinforced under a fixed ratio 1 schedule of cocaine reinforcement (0.15 mg of cocaine hydrochloride/50-µL infusion, delivered over 2.25 s, i.v.; NIDA Drug Supply Program, Research Triangle Park, NC) with a 20-s timeout period. Active-lever responses during timeouts and inactive-lever responses throughout the session were not reinforced. Training continued until the rats obtained at least 10 cocaine infusions per session on at least 10 days. Next, all rats received seven daily 2-h extinction training sessions in the alternate context. During extinction training, lever presses were not reinforced. After extinction session 4, rats were acclimated to the microinfusion procedure. Injection cannulae (33-Ga, Plastics One) were inserted 2 mm past the tip of the guide cannulae. The injection cannulae remained in place for 4 min but fluid was not infused.

### Experiment 1: Effects of post-retrieval AM251 administration in the BLA on drug context-induced cocaine seeking three days later

Twenty-four h after extinction session 7, rats were re-exposed to the cocaine-paired context for 15 min to trigger cocaine-memory retrieval and reconsolidation (**Fig. 1A**). During the session, cocaine reinforcement was withheld to prevent acute cocaine effects on neurotransmission and endocannabinoid mobilization, independent of memory destabilization [20-21]. Immediately after the session, rats received bilateral intra-BLA microinfusions of the CB1R antagonist/inverse agonist, *N*-(Piperidin-1-yl)-5-(4-iodophenyl)-1-(2,4-dichlorophenyl)-4-methyl-1*H*-pyrazole-3-carboxamide (**AM251**, 0.3 µg/0.5 µL/hemisphere; *n* = 9; Sigma Aldrich, St. Louis, MO), or vehicle (**VEH**; 8% DMSO, 5% Tween80 in saline; 0.5 µL/hemisphere; *n* = 11) over 2 min. This intra-BLA dose of AM251 is sufficient to impair contextual fear-memory reconsolidation [15]. After treatment, daily extinction-training sessions resumed in the extinction context until the rats reached the extinction criterion (i.e., ≤ 25 active-lever presses per session on two consecutive days). Non-reinforced lever responses were assessed during the first extinction session following treatment to evaluate possible off-target treatment effects on extinction memories. Twenty-four h after the last extinction session, cocaine-seeking behavior was assessed in the cocaine-paired context for 2 h.

**FIGURE 1.**
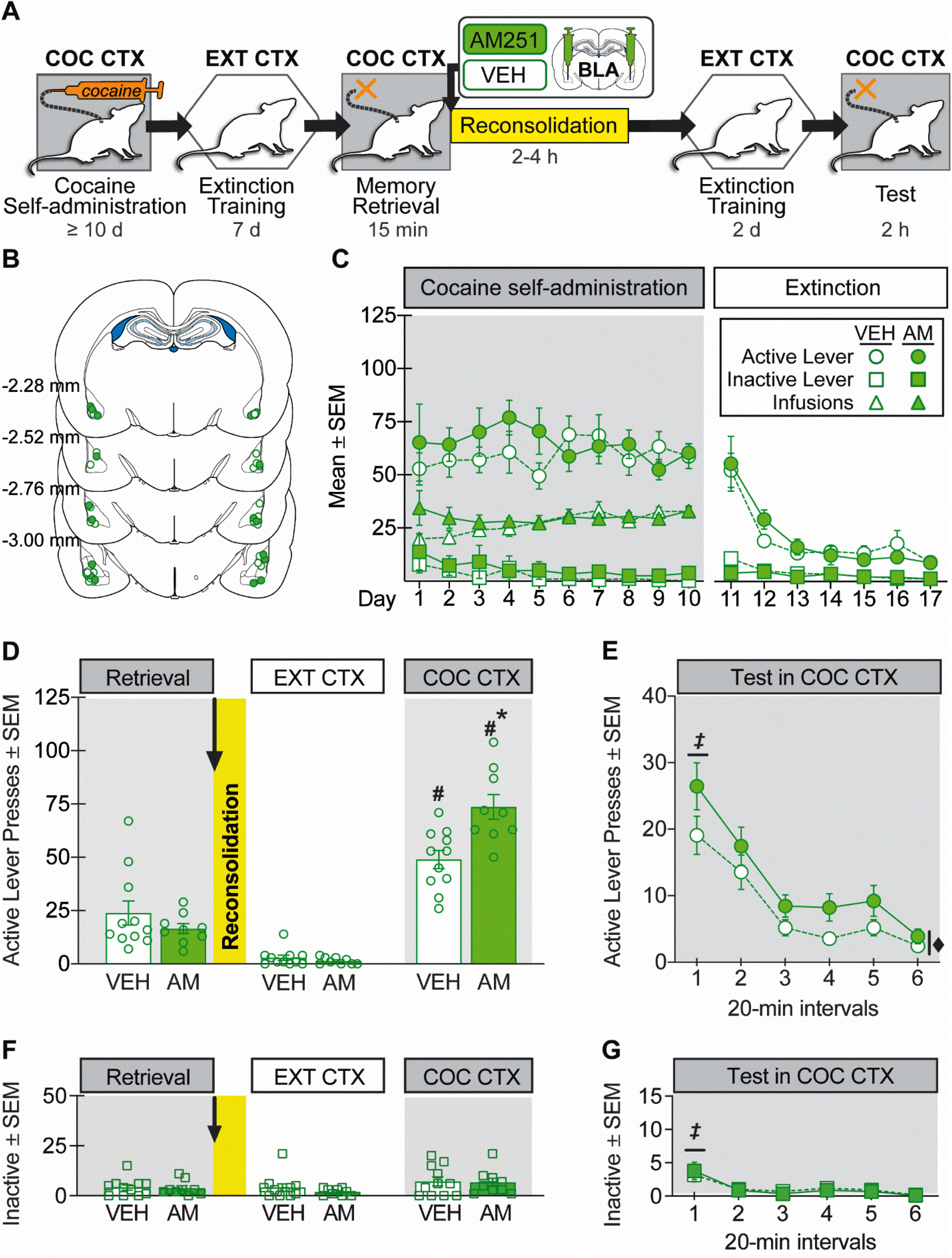
Intra-BLA AM251 administration *during* cocaine-memory reconsolidation increases drug context-induced cocaine seeking three days later. **(A)** Experimental timeline. After cocaine self-administration training in one context (**COC CTX**), and extinction training in a different context (**EXT CTX**), rats received bilateral intra-BLA administration of the CB1R antagonist, AM251 (**AM**; 0.3 µg/0.5µL per hemisphere; *n* = 9) or VEH (*n* = 11) immediately after the 15-min cocaine-memory retrieval session (**RETRIEVAL**). After two additional extinction sessions in the EXT CTX with ≤ 25 active lever responses, cocaine-seeking behavior was tested in the COC CTX. **(B)** Schematic of cannula placements. Symbols represent the most ventral point of injection cannula tracts for rats that received VEH (*open circles)* or AM251 *(closed circles)*. **(C)** Cocaine infusions and/or active- and inactive-lever responses (mean ± SEM) during cocaine self-administration (last 10 d) and extinction training prior to AM251 or VEH treatment. **(D)** Active-lever responses (mean ± SEM) at RETRIEVAL (before treatment) and upon first re-exposure to the EXT CTX and COC CTX after treatment. **(E)** Time course of active-lever responses (mean ± SEM) at test in the COC CTX. **(F)** Inactive-lever responses (mean ± SEM) during RETRIEVAL and upon first re-exposure to the EXT CTX and COC CTX. **(G)** Time course of inactive-lever responses (mean ± SEM) at test in the COC CTX. ***Symbols***: ANOVA ***#***context simple main effect, Sidak’s test, p < 0.05; ***\****treatment simple main effect, Sidak’s test, p < 0.05; ^*‡*^time simple main effect, Tukey’s tests, intervals 1 > 2-6, p < 0.05; ^♦^treatment main effect, p < 0.05.

### Experiment 2: Effects of delayed AM251 administration in the BLA on drug context-induced cocaine seeking three days later

Experiment 2 evaluated whether the intra-BLA AM251 effects observed in Experiment 1 required manipulation when memories were unstable (i.e., within 2-4 h after memory retrieval) prior to reconsolidation [22]. The procedures were identical to those in Experiment 1 except that rats received AM251 (*n* = 8) or VEH (*n* = 6) six h after the memory retrieval session (**Fig. 2A**).

**FIGURE 2.**
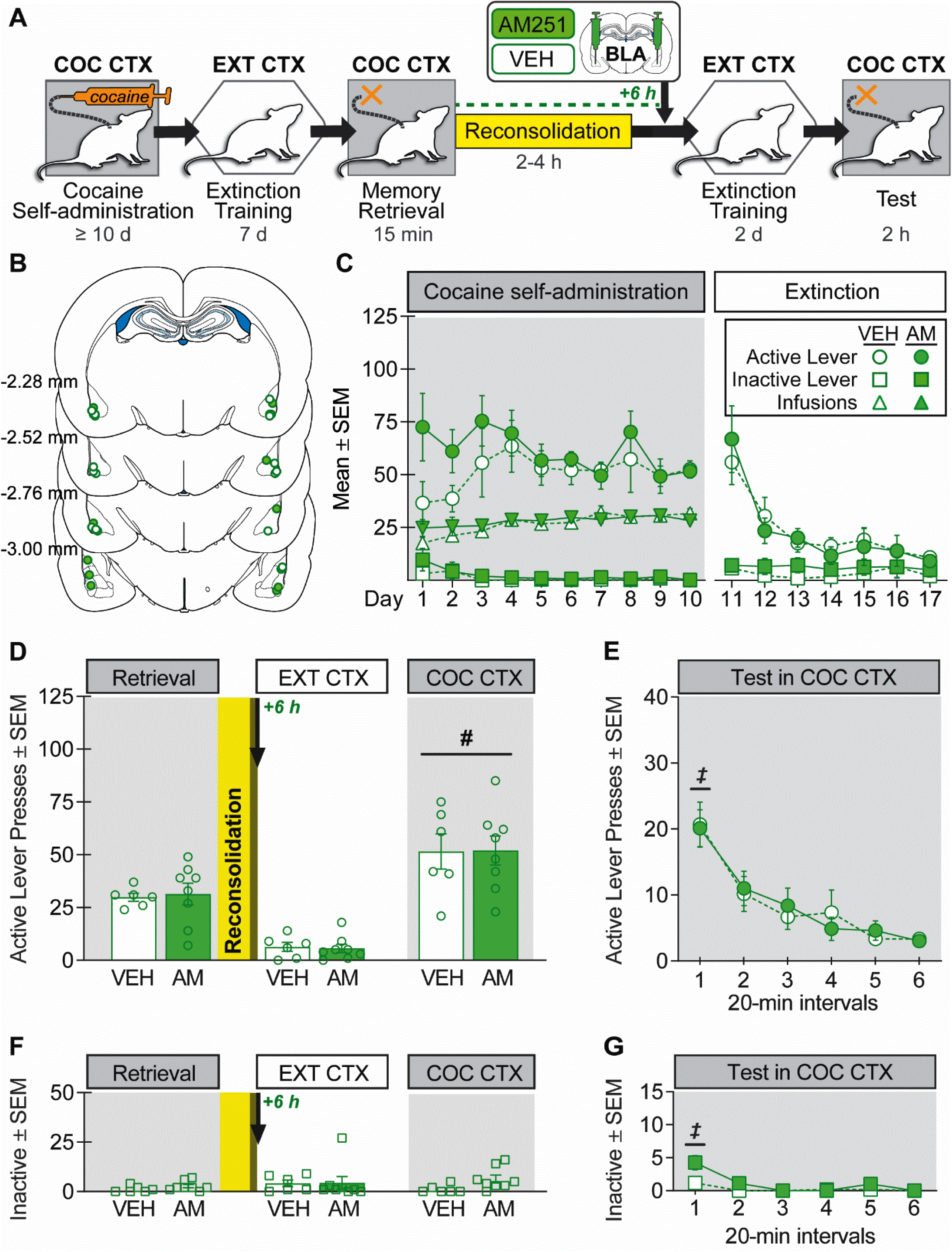
Intra-BLA AM251 administration *after* memory reconsolidation does not alter drug context-induced cocaine seeking three days later. **(A)** Experimental timeline. After cocaine self-administration training in one context (**COC CTX**) and extinction training in a different context (**EXT CTX**), rats received bilateral intra-BLA administration of the CB1R antagonist, AM251 (**AM**; 0.3 µg/0.5µL per hemisphere; *n* = 8) or VEH (*n* = 6) six hours after the 15-min cocaine-memory retrieval session (**RETRIEVAL**), after memory reconsolidation was completed. After two additional extinction sessions in the EXT CTX with ≤ 25 active lever responses, cocaine-seeking behavior was tested in the COC CTX. **(B)** Schematic of cannula placements. Symbols represent the most ventral point of injection cannula tracts for rats that received VEH (*open circles)* or AM251 *(closed circles)*. **(C)** Cocaine infusions and/or active- and inactive-lever responses (mean ± SEM) during cocaine self-administration (last 10 d) and extinction training prior to AM251 or VEH treatment. **(D)** Active-lever responses (mean ± SEM) at RETRIEVAL (before treatment) and upon first re-exposure to the EXT CTX and COC CTX after treatment. **(E)** Time course of active-lever responses (mean ± SEM) at test in the COC CTX. **(F)** Inactive-lever responses (mean ± SEM) during RETRIEVAL and upon first re-exposure to the EXT CTX and COC CTX. **(G)** Time course of inactive-lever responses (mean ± SEM) at test in the COC CTX. ***Symbols***: ANOVA, ***#***context main effect, p < 0.05 ^*‡*^time simple main effect, Tukey’s tests, intervals 1 > intervals 2-6, p < 0.05.

### Experiment 3: Effects of post-retrieval AM251 administration in the pCPu on drug context-induced cocaine seeking three days later

Experiment 3 evaluated whether the AM251 effects observed in Experiment 1 were anatomically specific to the BLA. The procedures were identical to those in Experiment 1 except that rats received AM251 (*n* = 9) or VEH (*n* = 7) into the pCPu immediately after the memory-retrieval session (**Fig. 3A**).

**FIGURE 3.**
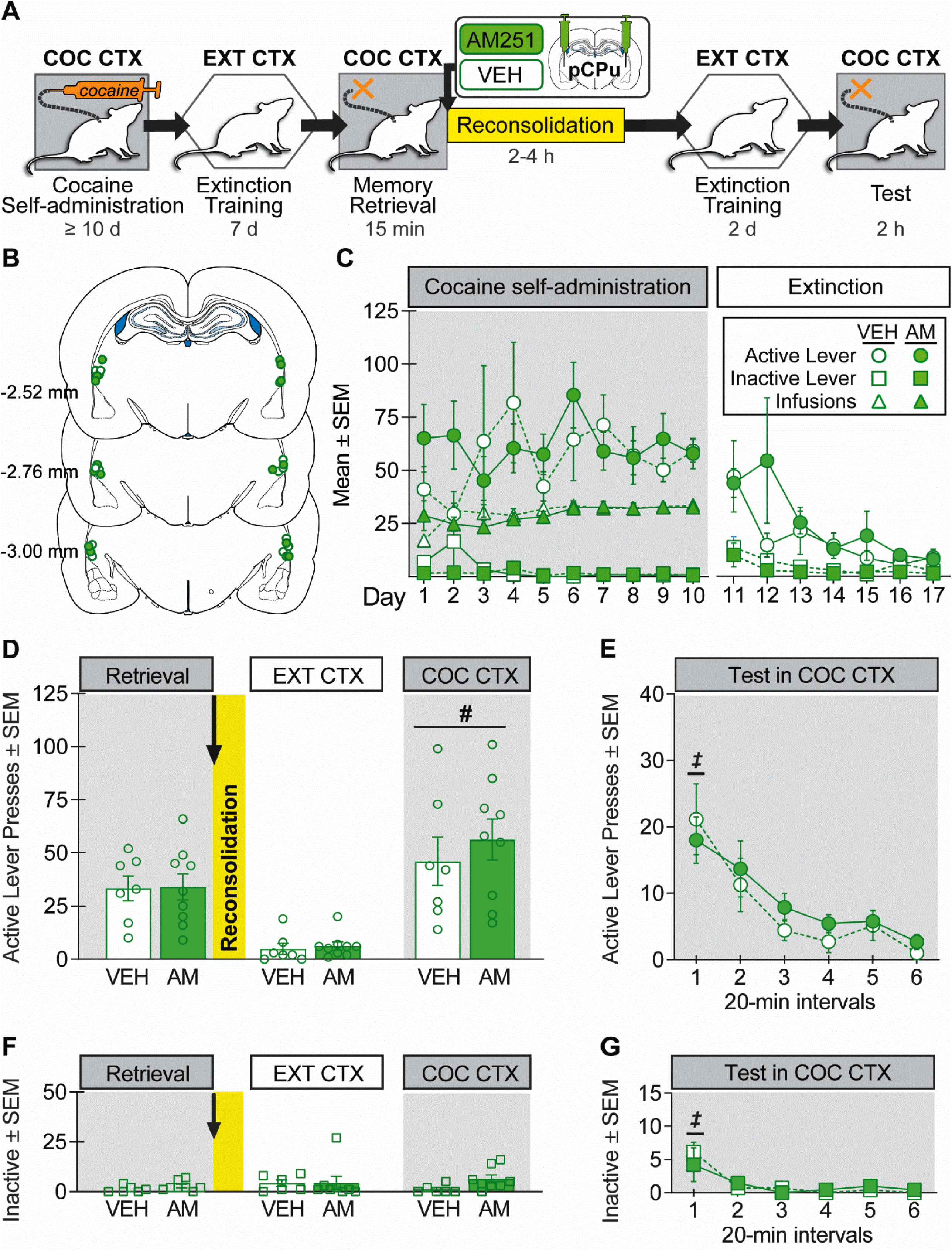
Intra-pCPu AM251 administration *during* memory reconsolidation does not alter drug context-induced cocaine seeking three days later. **(A)** Experimental timeline. After cocaine self-administration training in one context (**COC CTX**) and extinction training in a different context (**EXT CTX**), rats received bilateral intra-pCPu administration of the CB1R antagonist, AM251 (**AM**; 0.3 µg/0.5µL per hemisphere; *n* = 9) or VEH (*n* = 7) immediately after the 15-min cocaine-memory retrieval session (**RETRIEVAL**). After two additional extinction sessions in the EXT CTX with ≤ 25 active lever responses, cocaine-seeking behavior was tested in the COC CTX. **(B)** Schematic of cannula placements. Symbols represent the most ventral point of injection cannula tracts for rats that received VEH (*open circles)* or AM251 *(closed circles)*. **(C)** Cocaine infusions and/or active- and inactive-lever responses (mean ± SEM) during cocaine self-administration (last 10 d) and extinction training prior to AM251 or VEH treatment. **(D)** Active-lever responses (mean ± SEM) at RETRIEVAL (before treatment) and upon first re-exposure to the EXT CTX and COC CTX after treatment. **(E)** Time course of active-lever responses (mean ± SEM) at test in the COC CTX. **(F)** Inactive-lever responses (mean ± SEM) during RETRIEVAL and upon first re-exposure to the EXT CTX and COC CTX. **(G)** Time course of inactive-lever responses (mean ± SEM) at test in the COC CTX. ***Symbols***: ANOVA, ***#***context main effect, p < 0.5; ^*‡*^time simple main effect, Tukey’s tests, interval 1 > intervals 2-6, p < 0.05.

### Experiment 4: Effects of post-retrieval WIN 55,212-2 administration in the BLA on context-induced cocaine-seeking behavior three days later

Experiment 4 evaluated the effects of intra-BLA CB1R agonist administration on cocaine-memory reconsolidation. The procedures were identical to those in Experiment 1, except that rats received bilateral intra-BLA microinfusions of the nonselective CB1/CB2R agonist, WIN 55,212-2 (**WIN**, 0.5 µg/0.5-µL infusion/hemisphere; *n* = 11; Tocris Bioscience, Minneapolis, MN) or VEH (10% DMSO, 10% Tween80 in saline; 0.5 µL-infusion/hemisphere; *n* = 11) immediately after the memory-retrieval session (**Fig. 4A**). This intra-BLA dose of WIN enhances nicotine-conditioned place preference memory consolidation [23].

**FIGURE 4.**
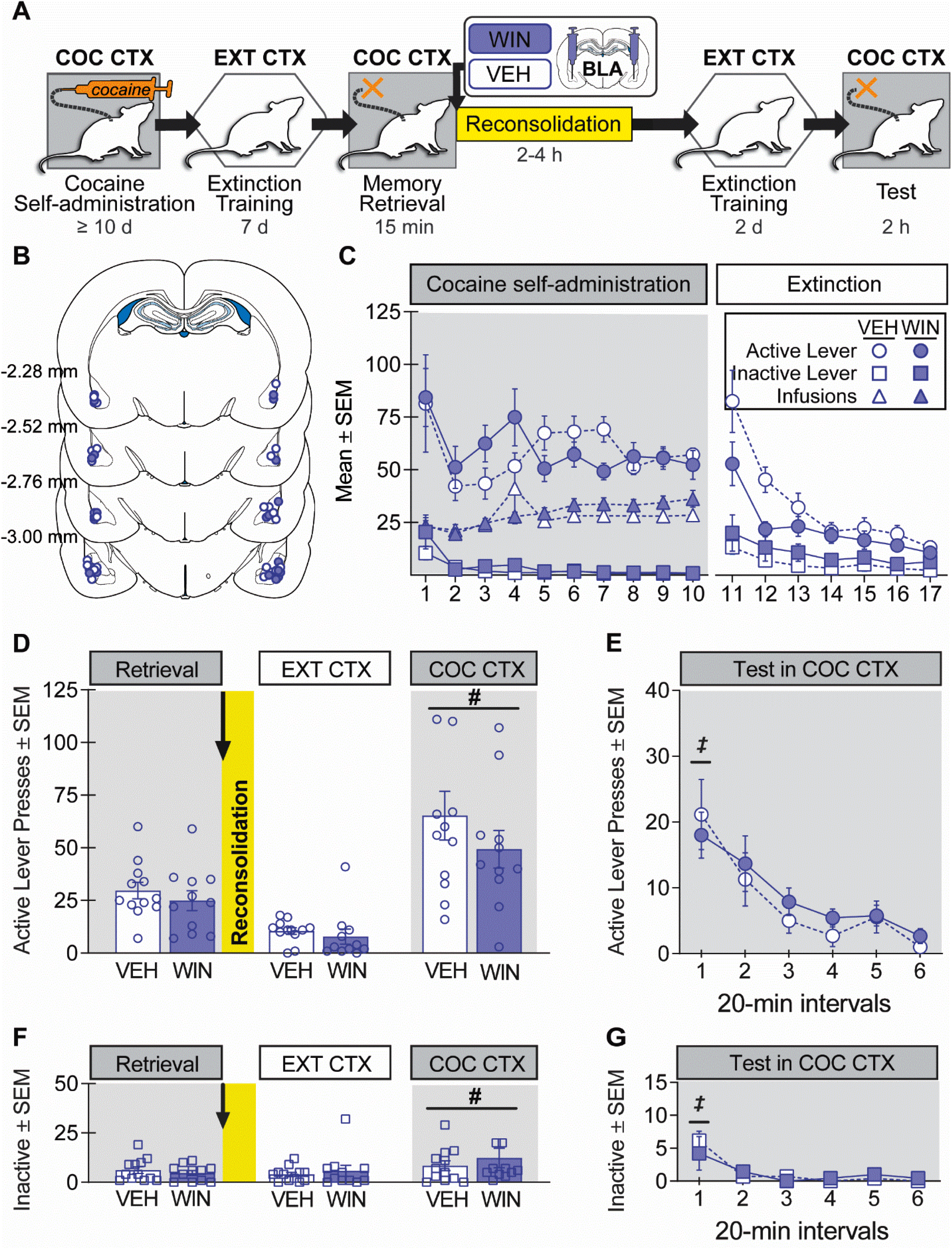
Intra-BLA WIN 55,212-2 administration *during* memory reconsolidation does not alter drug context-induced cocaine seeking three days later. **(A)** Experimental timeline. After cocaine self-administration training in one context (**COC CTX**) and extinction training in a different context (**EXT CTX**), rats received bilateral intra-BLA administration of the CB1R agonist, WIN 55,212-2 (**WIN**; 0.5 µg/0.5µL per hemisphere; *n* = 11) or VEH (*n* = 11) immediately after the 15-min cocaine-memory retrieval session (**RETRIEVAL**). After two additional extinction sessions in the EXT CTX with ≤ 25 active lever responses, cocaine-seeking behavior was tested in the COC CTX. **(B)** Schematic of cannula placements. Symbols represent the most ventral point of injection cannula tracts for rats that received VEH (*open circles)* or AM251 *(closed circles)*. **(C)** Cocaine infusions and/or active- and inactive-lever responses (mean ± SEM) during cocaine self-administration (last 10 d) and extinction training prior to AM251 or VEH treatment. **(D)** Active-lever responses (mean ± SEM) at RETRIEVAL (before treatment) and upon first re-exposure to the EXT CTX and COC CTX after treatment. **(E)** Time course of active-lever responses (mean ± SEM) at test in the COC CTX. **(F)** Inactive-lever responses (mean ± SEM) during RETRIEVAL and upon first re-exposure to the EXT CTX and COC CTX. **(G)** Time course of inactive-lever responses (mean ± SEM) at test in the COC CTX. ***Symbols***: ANOVA, ***#***context main effect, p < 0.05; ^*‡*^time simple main effect, Tukey’s tests, interval 1 > intervals 2-6, p < 0.05.

### Experiment 5: Effects of post-retrieval AM251 administration in the BLA on serum corticosterone concentrations during cocaine-memory reconsolidation

Experiment 5 examined the effects of cocaine-memory retrieval and intra-BLA AM251 treatment on serum corticosterone concentrations during the first 90 min of memory reconsolidation [16]. The training procedures were identical to those in Experiment 1, except that rats were acclimated to blood sample collection via tail nick (∼200 µL/sample) immediately before and after extinction session 6. Pre-session baseline blood samples were collected immediately before extinction session 7 (**Baseline**). Post-session blood samples were collected immediately after extinction session 7 (**Post-EXT**) and immediately after the memory-retrieval session in the cocaine-paired context (**Post-COC**; *n =* 11) or after comparable exposure to the home cage (**Post-Home**; *n* = 11) on post-cocaine day 8. AM251 (*n* = 6,6) or VEH (*n* = 5,5) was administered into the BLA immediately after the memory-retrieval session or exposure to the home cage. Post-treatment blood samples were collected 30, 60, and 90 min later (**Fig. 5A**). Blood samples were centrifuged at 4 °C. Blood serum was collected and stored at -20 °C. Samples were assayed in duplicates using the MP Biomedicals Corticosterone RIA kit for rats and mice (intra-assay coefficient of variation = 1.77 %, lower limit of detectability = 25 ng/mL).

**FIGURE 5.**
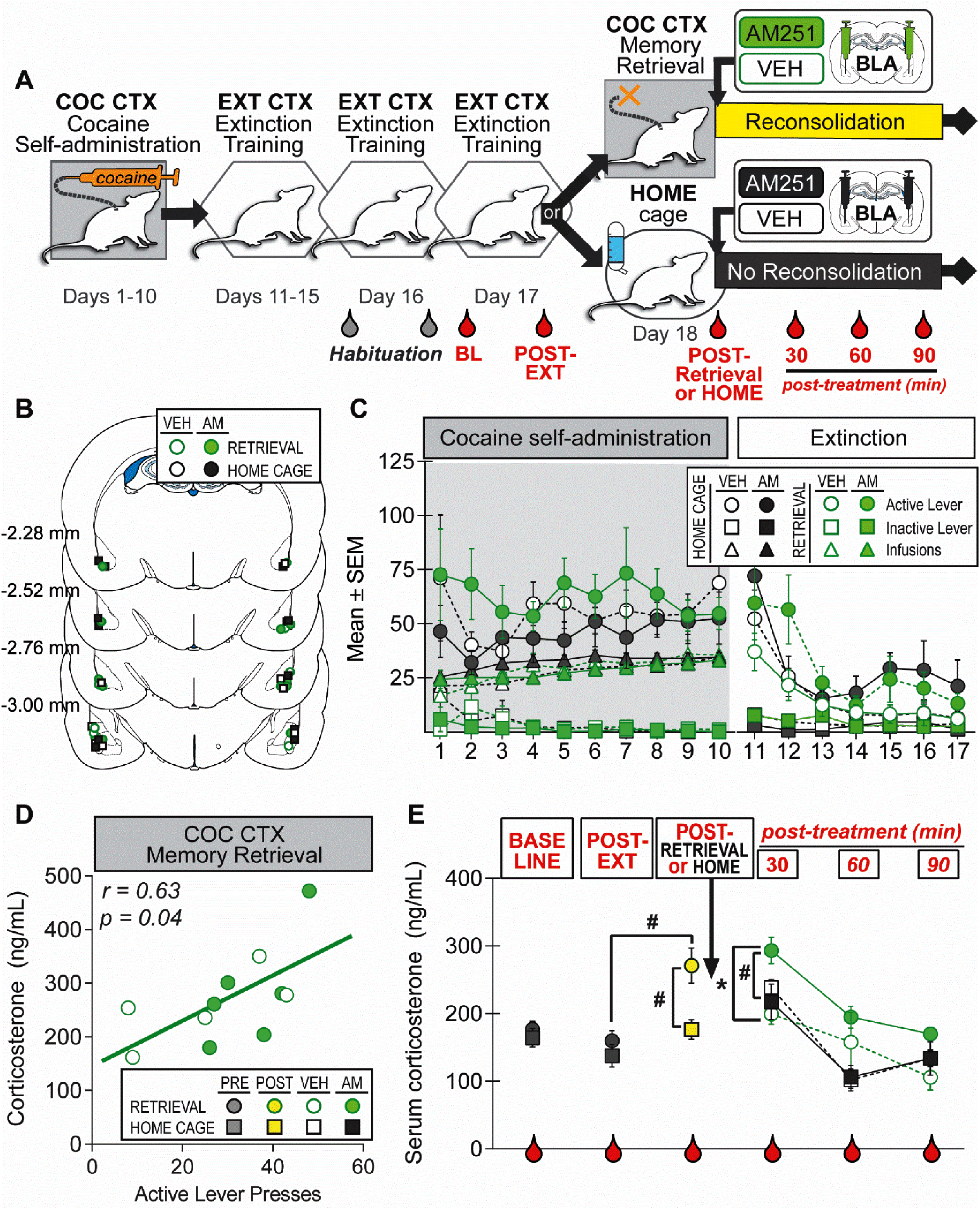
Intra-BLA AM251 administration prolongs memory retrieval-induced increase in blood serum corticosterone concentrations during cocaine-memory reconsolidation. **(A)** Experimental timeline. Rats received cocaine self-administration training in one context (**COC CTX**) and extinction training in a different context (**EXT CTX**). Rats were habituated to the tail-nick procedure before and after extinction session 6 (*gray symbols*). Blood samples (*red symbols*) were collected immediately prior to extinction session 7 (Baseline, **BL**), after extinction session 7 (**POST-EXT**), after either the 15-min cocaine memory-retrieval session (**POST-RETRIEVAL**; *n = 11*) or comparable exposure to the home cage (**POST-HOME**; *n = 11*), and after intra-BLA infusions of AM251 (**AM**; 0.3 µg/0.5µL per hemisphere) or VEH at 30-minute intervals (**30, 60, 90**). **(B)** Schematic of cannula placements in the BLA with symbols representing the most ventral point of injection cannula tracts for rats that were re-exposed to the COC CTX or home cage followed by VEH or AM251 treatment. **(C)** Cocaine infusions and/or active- and inactive-lever responses (mean ± SEM) during cocaine self-administration (last 10 d) and extinction training prior to memory retrieval and treatment manipulations. **(D)** Significant direct relationship between active-lever responses during the memory-retrieval session and POST-RETRIEVAL corticosterone concentrations before treatment (Pearson’s r). **(E)** Blood serum corticosterone concentrations (mean ± SEM) pre session (BASELINE), post session (POST-EXT, POST-EXT < POST-RETRIEVAL), and at 30, 60, and 90 min post treatment (AM251/retrieval > VEH/retrieval, AM251/home cage). ***Symbols***: ***#***one-way ANOVA, Tukey’s tests, p < 0.05; ***\****2 × 2 x 3 ANOVA, treatment simple main effects, Sidak’s tests, p < 0.05; ^*‡*^time simple main effect, Tukey’s tests, 30-min time point > 60-min and 90-min time points, p < 0.05.

### Histology

Rats were overdosed with a cocktail of ketamine and xylazine (300 and 15 mg/kg, respectively, i.p.). The brains were removed, flash frozen in isopentane, and stored at -80 °C. Forty-µm coronal brain sections were collected and stained with cresyl violet (Kodak, Rochester, NY, USA) to visualize cannula placement. Data of rats with cannula placements outside of the BLA or pCPu were excluded from the statistical analysis.

### Data analysis

Potential pre-existing group differences in cocaine intake and lever presses during (a) drug self-administration training (last 10 days), (b) extinction training (7 days), and (c) during the memory-retrieval session were analyzed using separate analyses of variance (ANOVA) with subsequent treatment group (VEH, AM251 or WIN) as the between-subjects factor and time (day) as the within-subject factor, as appropriate. Lever presses during the first post-treatment extinction session and the test session in the cocaine context were analyzed using ANOVAs with treatment (AM251 or WIN, VEH) as the between-subjects factor and context (extinction, cocaine-paired) and time (20-min interval) as within-subjects factors, where appropriate. Serum corticosterone concentrations were analyzed using ANOVAs with context (Post-EXT, Post-COC, Post-Home), memory retrieval (memory retrieval, no-memory retrieval), and treatment (AM251, VEH) as between-subjects factors, and time or context (Baseline, Post-EXT or Post-COC; and 30-min intervals) as within-subjects factors, where appropriate. Significant interaction and main effects were further analyzed using *post hoc* Sidak’s multiple comparisons tests or Tukey’s tests, where appropriate. Alpha was set at 0.05 for all analyses.

## RESULTS

### Behavioral History

The groups did not differ in drug intake during cocaine self-administration training or in active-or inactive-lever responding during cocaine self-administration training, extinction training, or cocaine-memory retrieval in Experiments 1-5 (see ANOVA results in ***Table S2***). The groups also did not differ in the mean number of sessions required to reach the extinction criterion prior to testing in Experiments 1-4 (mean ± SEM = 2.16 ± 0.14). Thus, testing in the cocaine-paired context occurred for most rats three days post treatment.

### Experiment 1: Intra-BLA AM251 administration during cocaine-memory reconsolidation increased subsequent drug context-induced cocaine seeking

Intra-BLA AM251 treatment immediately after cocaine-memory retrieval selectively increased drug context-induced cocaine-seeking behavior three days later relative to VEH (ANOVA context x treatment interaction, F_(1,18)_ = 14.60, p = 0.001; context main effect F_(1,18)_ = 299.20, p < 0.0001; treatment main effect, F_(1,18)_ = 9.79, p = 0.006). Active-lever responding was higher in the cocaine-paired context than in the extinction context (**Fig. 1D;** Sidak’s test, p < 0.05). Furthermore, AM251 administered after memory retrieval increased responding in the cocaine-paired context (Sidak’s test, p < 0.05), but not in the extinction context, relative to VEH. Time-course analysis indicated that active-lever responding declined over time in the cocaine-paired context at test (ANOVA time main effect, F_(5,90)_ = 25.61, p < 0.0001, Tukey’s tests, interval 1 > 2-6, p < 0.05; time x treatment interaction, F_(5,90)_ = 0.44, p = 0.82), and AM251 increased responding relative to VEH independent of time (**Fig. 1E**; treatment main effect, F_(1,18)_ = 12.46, p = 0.002). Inactive-lever responding remained low in both contexts (**Fig.1F**; all Fs ≤ 3.96, ps ≥ 0.06), and it declined during the test session independent of treatment (**Fig. 1G**; ANOVA time main effect only, F_(5,90)_ = 10.45, p < 0.0001, Tukey test, interval 1 > 2-6, p < 0.05; all other Fs ≤ 0.23, ps ≥ 0.87).

### Experiment 2: Intra-BLA AM251 administration after memory re-stabilization did not alter subsequent drug context-induced cocaine seeking

Intra-BLA AM251 administration six h after memory retrieval did not alter cocaine-seeking behavior three days later relative to VEH (**Fig. 2D**; ANOVA treatment main and interaction effects, Fs ≤ 0.01, ps ≥ 0.90). Thus, active-lever responding was higher in the cocaine-paired context than in the extinction context, independent of treatment (context main effect, F_(1,12)_ = 86.26, p < 0.0001). Time-course analysis confirmed that active-lever responding declined over time in the cocaine-paired context at test, independent of treatment (**Fig. 2E**; ANOVA time main effect, F_(5,60)_ = 18.02, p < 0.0001, Tukey test, interval 1 > 2-6, p < 0.05; all other Fs ≤ 0.001, ps ≥ 0.93). Inactive-lever responding remained low in both contexts (**Fig.2F**; all Fs ≤ 1.51, ps ≥ 0.24), and it declined during the test session independent of treatment (**Fig. 2G**; ANOVA time main effect F_(5,60)_ = 18.02, p < 0.0001, Tukey test, interval 1 > 2-6, p < 0.05; all other Fs ≤ 0.26, ps ≥ 0.93).

### Experiment 3: Intra-pCPu AM251 administration during cocaine-memory reconsolidation did not alter subsequent drug context-induced cocaine seeking

Intra-pCPu AM251 administration immediately after cocaine-memory retrieval did not alter cocaine-seeking behavior relative to VEH three days later (**Fig. 3D**; ANOVA, all Fs_(1,14)_ ≤ 0.70, ps ≥ 0.42). Active-lever responding was higher in the cocaine-paired context then in the extinction context, independent of treatment (context main effect, F_(1,14)_ = 38.59, p < 0.0001). Furthermore, time-course analysis indicated that responding declined over time in the cocaine-paired context, independent of treatment (**Fig. 3E**; ANOVA, time main effect, F_(5,70)_ = 15.75, p < 0.0001, Tukey test, interval 1 > 2-6, p < 0.05; all other Fs ≤ 0.49, ps ≥ 0.68). Inactive-lever responding remained low in both contexts (**Fig.1F**; all Fs ≤ 2.28, ps ≥ 0.15), and it declined during the test session independent of treatment (**Fig. 1G**; ANOVA time main effect F_(5,70)_ = 10.35, p < 0.004, Tukey test, interval 1 > 2-6, p < 0.05; all other Fs ≤ 0.75, ps ≥ 0.588).

### Experiment 4: Intra-BLA WIN 55,212-2 administration during memory reconsolidation failed to alter subsequent drug context-induced cocaine seeking

Intra-BLA WIN administration immediately after cocaine-memory retrieval did not alter cocaine-seeking behavior three days later (**Fig. 4D**; ANOVA treatment main and interaction effects, Fs ≤ 1.75, ps ≥ 0.20). Active-lever responding was higher in the cocaine-paired context than in the extinction context, independent of treatment (context main effect, F_(1,21)_ = 41.35, p < 0.0001). Similarly, time-course analysis indicated that active-lever responding declined over time at test, independent of treatment (**Fig. 4E**; ANOVA time main effect, F_(5,105)_ = 33.38, p < 0.0001, Tukey test, interval 1 > 2-6, p < 0.05; all other Fs ≤ 1.30, ps ≥ 0.27). Inactive-lever responding was higher in the cocaine-paired context than in the extinction context, independent of treatment (**Fig. 4F**; ANOVA context main effect, F_(1,21)_ = 5.47, p = 0.03; all other Fs_(1,21)_ ≤ 1.18, ps ≥ 0.29), and it declined during the test session, independent of treatment (**Fig. 4G;** ANOVA time main effect, F_(5,105)_ = 21.52, p < 0.0001, Tukey test, interval 1 > 2-6, p < 0.05; all other Fs ≤ 0.72, ps ≥ 0.41).

### Experiment 5: Intra-BLA AM251 administration potentiated increases in serum corticosterone concentrations during cocaine-memory reconsolidation

There were no pre-existing differences between the groups in baseline pre-session and post-extinction session serum corticosterone concentrations (ANOVA, all Fs ≤ 2.95, ps ≥ 0.10). Cocaine-memory retrieval (i.e., cocaine-paired context re-exposure) increased corticosterone concentrations compared to no-memory retrieval (i.e., extinction context or home cage re-exposure; **Fig. 5E**; ANOVA, F_(3,40)_ = 10.01, p < 0.0001; Tukey’s tests, p < 0.05). Furthermore, active-lever responses during the cocaine-memory retrieval session, but not during the non-memory retrieval session (extinction, not shown), positively correlated with corticosterone concentrations immediately post session (**Fig. 5D**; Pearson’s r = 0.63, p = 0.04).

Following intra-BLA AM251 or VEH treatment, serum corticosterone concentrations declined over time (**Fig. 5E**: 2×2×3 ANOVA time main effect, F_(2,36)_ = 43.51, p < 0.0001, Tukey’s tests, 30-min time point > 60-min and 90-min time points, p < 0.05). However, intra-BLA AM251 administered after memory retrieval resulted in higher serum corticosterone concentrations relative to VEH after memory retrieval and relative to AM251 after no-memory retrieval (2×2×3 ANOVA treatment x retrieval interaction, F_(1,18)_ = 5.266, p = 0.03, Sidak’s tests, p < 0.05; all other Fs ≤ 4.21, ps ≥ 0.06).

## DISCUSSION

In the present study, we used site-specific pharmacological manipulations to examine the contribution of BLA CB1R populations to cocaine-memory reconsolidation for the first time in an instrumental rat model of drug relapse. Our findings indicate that CB1R signaling in the BLA limits the strength of reconsolidating cocaine memories in a time-dependent manner, possibly by reducing the impact of memory retrieval-induced HPA axis activation on neural circuits engaged during memory reconsolidation.

BLA CB1R antagonism by AM251 immediately after cocaine-memory retrieval (i.e., during memory reconsolidation) increased subsequent drug context-induced cocaine-seeking behavior (**Fig. 1**). This effect might reflect AM251-induced augmentation of (a) memory re-stabilization efficiency or (b) memory strength itself at the time of reconsolidation, as opposed to protracted enhancement of the performance of cocaine-seeking behavior, since AM251 treatment six h after cocaine-memory retrieval did not facilitate this behavior at test, relative to VEH (**Fig. 2**). The memory retrieval-dependent effects of AM251 were anatomically specific to the BLA, as AM251 infusions into the dorsally adjacent pCPu after cocaine-memory retrieval failed to alter subsequent cocaine-seeking behavior relative to VEH (**Fig. 3**). Together, these findings provide the first demonstration that BLA CB1R signaling may play a negative regulatory role in appetitive-memory reconsolidation. These findings expand upon a seemingly inconsistent literature indicating that CB1R antagonist [15] or agonist [14] treatments in the BLA impair fear-memory reconsolidation. Notably, systemic administration of the same CB1R agonist can either facilitate or impair object-recognition memory *consolidation* depending on whether conditioning takes place in an unhabituated (thus emotionally arousing) or a familiar environment [24], and similar mechanisms may be at play in memory *reconsolidation*. Accordingly, varying results may reflect that specific appetitive and aversive emotional states and arousal evoked in these paradigms may differently alter endocannabinoid recruitment and, thus, the functional contribution of CB1R populations to memory reconsolidation [24-26].

Intra-BLA administration of the non-selective CB1R agonist, WIN, after memory retrieval failed to alter subsequent drug context-induced cocaine-seeking behavior (**Fig. 4**); even though BLA CB1R antagonism potentiated this behavior. It is unlikely that the lack of a WIN effect reflected insufficient dosing, since this intra-BLA dose (0.5 µg/hemisphere) inhibits nicotine-memory consolidation [23] while even higher doses of WIN (1-5 µg/hemisphere) fail to have consistent effects on fear-memory reconsolidation [14, 27]. Therefore, it is possible that BLA CB1R signaling is insufficient, but necessary, for limiting cocaine-memory strength during reconsolidation *per se*. Alternatively, we propose that nonselective effects of WIN interfered with our ability to selectively increase BLA CB1R signaling relevant for cocaine-memory reconsolidation. Unlike AM251, which exerts selective effects on BLA cell populations that are experiencing dynamic endocannabinoid mobilization, WIN stimulates CB1Rs on both glutamatergic and GABAergic terminals within the BLA, likely with opposing effects on reconsolidation. Additionally, WIN is a nonselective agonist with 19-fold greater selectivity for CB2Rs than for CB1Rs [28], both of which are expressed in the BLA [29]. Finally, WIN is a biased CB1R agonist that exhibits lower efficacy to stimulate Gi/o- and Gq-coupled CB1Rs than the endocannabinoids, anandamide (**AEA**) and 2-arachydonoylglicerol (**2-AG**), but higher efficacy to stimulate arrestin-2-coupled CB1Rs than AEA [30]. In conclusion, differential recruitment of distinct CB1R- and CB2R-bearing cell populations and CB1Rs with different effector systems, may contribute to the inconsistencies between the effects of WIN and AM251 in the present study as well as to the discrepancies in the effects of WIN across various fear-memory reconsolidation paradigms.

It has been well documented that exposure to drug-associated stimuli triggers HPA axis activation in cocaine users [31] and cocaine-trained rats [16, 32]. Furthermore, stress-induced reductions in endocannabinoid tone in the BLA facilitate HPA axis activation [17, 33]. In the present study, drug context-induced cocaine-memory retrieval resulted in a significant increase in serum corticosterone concentrations compared to two control conditions: re-exposure to the extinction context or the home cage (**Fig. 5**). Furthermore, there was a direct relationship between serum corticosterone concentrations and cocaine-seeking behavior during the memory retrieval session (**Fig. 5**). The magnitude of the corticosterone response was comparable to those observed upon exposure to cocaine-paired contextual stimuli [32] or mild stressors [34], such as elevated platform stress [35] and restraint stress [36], in previous studies.

Remarkably, intra-BLA AM251 administration prolonged the drug context-induced corticosterone response during memory reconsolidation (i.e., after cocaine-memory retrieval) relative to VEH (**Fig. 5**), while it did not alter corticosterone secretion following no-memory retrieval. These findings are consistent with extant literature indicating that intra-BLA AM251 treatment alone is not anxiogenic [37], and it selectively enhances stress-induced, but not baseline, serum corticosterone concentrations [17]. Therefore, in the present study, intra-BLA AM251 administration might prolong a memory retrieval-induced arousal state that increased the strength of reconsolidating cocaine memories or the efficiency of memory reconsolidation. Accordingly, BLA CB1R signaling may gate memory strength and protect against the development of maladaptively strong and intrusive cocaine memories.

The contributions of specific endocannabinoids, including AEA and 2-AG, to CB1R-mediated effects on cocaine-memory reconsolidation have yet to be determined, but some insights may be gained from the stress literature. Upon exposure to a stressor, *AEA tone diminishes* in the BLA, due to corticotropin-releasing factor (**CRF**)-induced stimulation of AEA hydrolysis [38]. This leads to delayed, *phasic 2-AG release* due to the disinhibition of BLA glutamatergic principal neurons [39-40] and thus metabotropic glutamate receptor-mediated, as well as glucocorticoid receptor-mediated, stimulation of 2-AG synthesis [40-42]. Similar to stressors, cocaine-memory retrieval stimulates HPA axis activity [16] (**Fig. 5**), and it likely reduces AEA levels and increases 2-AG levels in the BLA during reconsolidation. As a CB1R antagonist, AM251 in the BLA inhibits 2-AG and AEA signaling and, as such, augments the impact of cocaine-memory retrieval on HPA axis activity, as indicated by the potentiated corticosterone response (**Fig. 5**). The resulting increase in BLA principal neuronal activity during memory reconsolidation may enhance cocaine-memory strength or storage efficiency, similar to fear memory consolidation [43]. Future studies will need to determine whether memory retrieval-induced alterations in BLA AEA, 2-AG, or both are critical for this phenomenon.

## CONCLUSIONS

While intra-BLA AM251 administration *enhanced*, systemic AM251 administration in our previous study *impaired* [7], cocaine-memory strength or reconsolidation efficiency. These findings indicate that functionally heterogeneous CB1Rs populations bidirectionally regulate cocaine-memory strength or reconsolidation as components of larger neural circuits. Although intra-BLA AM251 prolonged the memory retrieval-induced increase in blood serum corticosterone concentrations, it is unlikely that corticosterone mediated the effects on memory strength within the BLA, because we have previously shown that intra-BLA glucocorticoid receptor antagonism *enhances* cocaine-memory reconsolidation [16]. Instead, CRF and/or norepinephrine may mediate the effects of BLA AM251 on memory reconsolidation, in the course of HPA axis stimulation. In support of this alternative, stressors elicit an increase in CRF immunoreactivity [44] and norepinephrine release in the BLA [45-46]. Furthermore, intra-BLA CRH receptor type 1 (Fuchs and Ritchie, unpublished) or ß-adrenergic receptor antagonism disrupts cocaine-memory reconsolidation ([47]; Fuchs and Higginbotham, unpublished).

Based on the emerging role of CB1Rs in memory reconsolidation, CB1R genetic polymorphisms and other factors that lead to abnormalities in CB1R signaling may regulate an individual’s susceptibility to SUDs and other psychiatric disorders that are characterized by pathologically strong maladaptive memories. Moreover, dysfunction of endocannabinoid recruitment upon exposure to drug-associated environmental stimuli and during subsequent drug-memory retrieval and reconsolidation may influence subsequent drug-relapse propensity. Interfering with neural mechanisms that enhance cocaine-memory strength during reconsolidation may be a useful adjunct to other approaches for drug-relapse prevention.

## Acknowledgments

The authors are grateful to Shi Min Tan and Ethan Hansen for excellent technical assistance as well as to Dr. Anthony Berger for help with the corticosterone assays.

## Author Contributions

JAH and RAF developed the study concept and experimental design; JAH, NMJ, RJM, and RAF collected the data; JAH, RAF, and RJM analyzed and interpreted the data; JAH and RAF wrote the manuscript with input from all authors. All authors have approved the version of the manuscript submitted for publication.

## SUPPLEMENTARY MATERIALS

**Table S1.**

| Context | Visual | Auditory | Olfactory | Tactile |
| --- | --- | --- | --- | --- |
| **A** | Continuous red house light | Intermittent tone (78 dB, 10 hz) | Vanilla-scented air freshener | Wire mesh flooring |
| **B** | Flashing white light above inactive lever | Continuous tone (78 dB, 2kHz) | Pine-scented air freshener | Slanted tile bisecting a steel grid flooring |

**Table S2.**
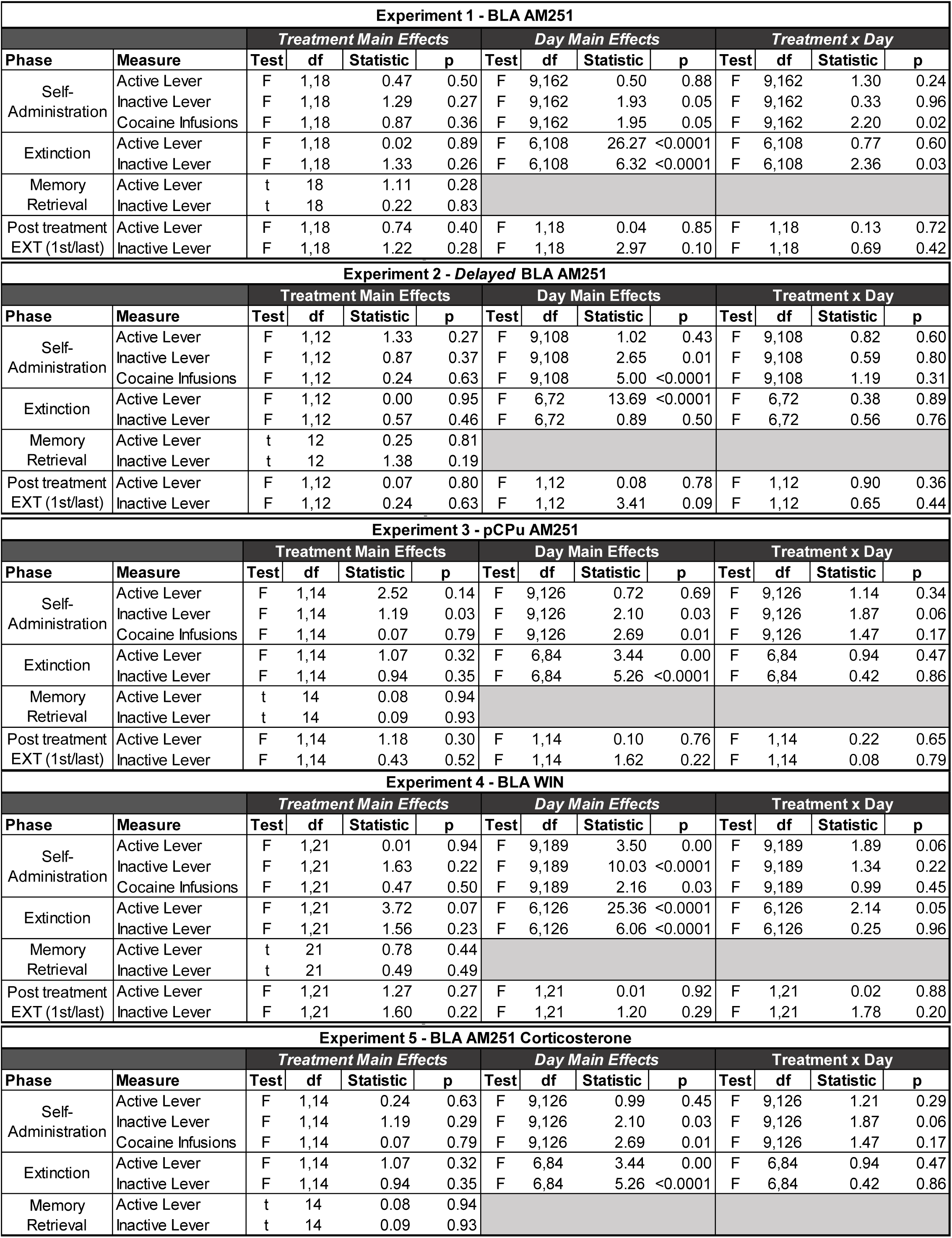
Behavioral History of Rats in Experiments 1-5.

